# An atlas of genetic variation for linking pathogen-induced cellular traits to human disease

**DOI:** 10.1101/202325

**Authors:** Liuyang Wang, Kelly J. Pittman, Jeffrey R. Barker, Raul E. Salinas, Ian B. Stanaway, Graham D. Williams, Robert J. Carroll, Tom Balmat, Andy Ingham, Anusha M. Gopalakrishnan, Kyle D. Gibbs, Alejandro L. Antonia, The eMERGE Network, Joseph Heitman, Soo Chan Lee, Gail P. Jarvick, Joshua C. Denny, Stacy M. Horner, Mark R. Delong, Raphael H. Valdivia, David R. Crosslin, Dennis C. Ko

## Abstract

Genome-wide association studies (GWAS) have identified thousands of genetic variants associated with disease. To facilitate moving from associations to disease mechanisms, we leveraged the role of pathogens in shaping human evolution with the Hi-HOST Phenome Project (H2P2): a catalog of cellular GWAS comprised of 79 phenotypes in response to 8 pathogens in 528 lymphoblastoid cell lines. Seventeen loci surpass genome-wide significance (p<5×10^−8^) for phenotypes ranging from pathogen replication to cytokine production. Combining H2P2 with clinical association data from the eMERGE Network and experimental validation revealed evidence for mechanisms of action and connections with diseases. We identified a SNP near *CXCL10* as a cis-cytokine-QTL and a new risk factor for inflammatory bowel disease. A SNP in *ZBTB20* demonstrated pleiotropy, partially mediated through NF*κ*B signaling, and was associated with viral hepatitis. Data are available in an H2P2 web portal to facilitate further interpreting human genome variation through the lens of cell biology.

## Introduction

The human genome has been shaped by migration, drift, admixture, and natural selection (*1–3*). One of the strongest driving forces in natural selection has been pathogens (*4, 5*), as first exemplified with A. C. Allison’s demonstration that sickle cell allele (rs334) conferred resistance to malaria (*6*). Red blood cells from individuals with this allele are resistant to *Plasmodium* infection (*7*). Similarly, human resistance to HIV infections afforded by the *CCR5Δ32* allele can also be seen at the level of individual T cells (*8–10*). Therefore, identification and characterization of human genetic differences that impact cellular traits can help mechanistically link human genetic variation to disease susceptibility.

Previous studies have examined the genetic basis of molecular traits in human populations. Expression quantitative trait loci (eQTL) studies in lymphoblastoid cell lines (LCLs) defined abundant associations between human SNPs and expression levels of nearby genes (*11, 12*). LCLs are EBV-transformed B cells that are highly similar to antigen-activated primary B cells (*13*). LCLs serve as a standardized resource for functional human genetic variation studies, as they have been densely genotyped by the International HapMap Project (*14, 15*) and the 1000 Genomes Project (*16*). As eQTLs are often shared across tissues (e.g. 88% of cis-eQTLs are shared among LCLs, fibroblasts, and primary T cells (*17*)), LCL eQTL studies have led to important insights not only in immunity-related diseases but also for disorders where B cells are not believed to be primary drivers of disease (*18*).

Using LCLs, we developed Hi-HOST (High-throughput Human in vitrO Susceptibility Testing) to identify human genetic differences in pathogen-induced cellular traits, serving as a cell biological link between eQTL studies and GWAS of human disease (*19, 20*). Hi-HOST uses live pathogens to examine variation in innate immune recognition, but also in cell biological and signaling processes that can be quantified as phenotypes for genome-wide association. This work therefore builds on a long tradition of using cellular microbiology to elucidate basic cell biology (*21*) and expands that utility to interpret the human genome. Using Hi-HOST, we leveraged LCL responses to *Salmonella enterica* to demonstrate that genetic variation in the methionine salvage pathway regulates pyroptosis, as well as human susceptibility to sepsis (*22, 23*). Similarly, we recently reported that a genetic variant in *VAC14* is associated with both increased *S.* Typhi invasion into LCLs and risk of typhoid fever in a Vietnamese population (*24*).

Here, we present the Hi-HOST Phenome Project (H2P2) to globally explore the genetic basis of cellular outcomes in response to infectious agents. Using 7 microbes and 1 bacterial toxin, we carried out GWAS of 79 host-pathogen phenotypes that serve as cellular readouts for processes such as endocytosis, endosomal trafficking, signal transduction, cell death, and transcriptional regulation. We identified 17 loci that reached the generally accepted threshold for genome-wide significance (p<5×10^−8^). We integrated H2P2 data with experimental validation and disease association data from the eMERGE Network PheWAS pipeline (*25*) to define new functions for genes in disease and provide new clues to disease pathophysiology.

## Results

### Phenotypic variation in H2P2 traits reveals biologically meaningful clusters

We measured variation in cellular traits in 528 LCLs stimulated with 7 different microbes and 1 bacterial toxin (Figure 1A). These LCLs have been genotyped (*16*) and consist of 4 human populations: ESN (Esan in Nigeria), GWD (Gambians in Western Divisions in the Gambia), IBS (Iberian Population in Spain), and KHV (Kinh in Ho Chi Minh City, Vietnam). The LCLs used in this study were all from parent-offspring trios, allowing for heritability estimation by parent-offspring regression and for carrying out combined GWAS analysis with protection from stratification through family-based association methods (*26, 27*).

**Figure 1.**
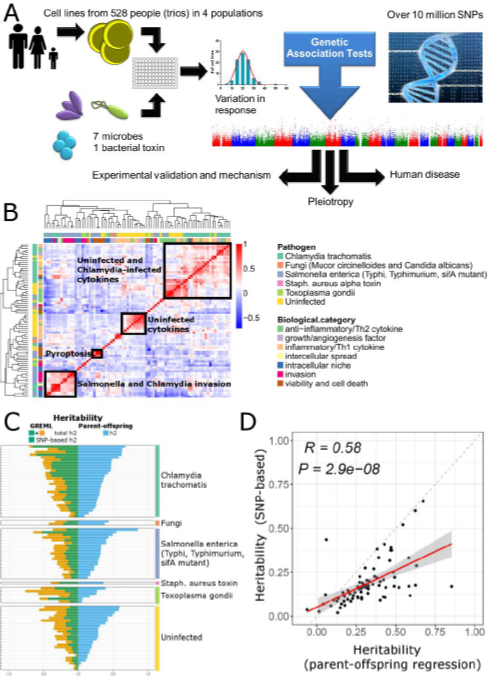
Inter-individual variation in H2P2 traits reveals clustering of phenotypes and heritable variation in cellular responses to infection. (A) Diagram of the H2P2 workflow for connecting genetic variation to cell biology. Lymphoblastoid cell lines (LCLs) from 528 people (parent-offspring trios) from 4 populations were exposed to 8 different stimuli for 79 phenotypes. Over 10 million SNPs were tested for association with each phenotype using family-based association implemented in PLINK (*26, 27*). (B) Clustering of H2P2 phenotypes reveals a map of trait similarity. Heatmap and dendrograms of hierarchical clustering based on inter-individual phenotypic variation. Spearman correlation is color-coded in the heatmap from blue (negative correlation) to red (positive correlation). Phenotypes are color-coded by stimuli (outer band) and biological category of phenotype (inner band). Several clusters of related traits are highlighted. (C) Narrow-sense heritability (h^2^) estimates for H2P2 phenotypes based on GREML vs. parent-offspring regression. h^2^ was estimated using genome-wide SNP data using the Zaitlen method of GREML for family data in GCTA or by parent-offspring regression. Both methods utilized a covariate for batch effects. The GREML method gives a SNP-based h^2^ (green) as well as a total h^2^ (yellow; the sum of SNP-based h^2^ (green) plus the non-SNP-based h^2^). Both methods indicate a large range of h^2^ estimates for different H2P2 traits and are consistent with many traits having a significant genetic component. (D) h^2^ estimates from parent-offspring regression vs. GREML SNP-based h^2^ are well correlated. Linear regression for all 79 H2P2 traits demonstrate the two methods give similar h^2^ estimates.

The microbes and toxin we selected affect billions of people worldwide. Non-typhoidal *Salmonella* infections caused 150 million diarrheal illnesses (gastroenteritis) and 0.6 million cases of invasive enteric disease (bacteremia) in 2010 (*28, 29*). Approximately 20 million cases of typhoid fever are caused by *Salmonella enterica* serovar Typhi (S. Typhi) every year (*30*). *Chlamydia trachomatis*, the most common bacterial sexually transmitted infection in the world, causes 100 million cases of genital tract infection every year (*31*), and 1.3 million individuals are blind due to ocular infection with *C. trachomatis* (*32*). *Staphylococcus aureus* is a common cause of skin and soft tissue infections, bacteremia, and infective endocarditis, and its alpha toxin, which is utilized in H2P2, is a key virulence determinant (*33*). *Candida albicans* and related fungal species are a frequent cause of genitourinary tract infection that can cause more severe disease in immunocompromised individuals and are the fifth most common cause of hospital-associated infections (*34*). *Mucor circinelloides* is another fungal species that causes severe infections, mucormycosis, in immunocompromised individuals (reviewed in (*35*)) and has also recently been connected to food-borne infections (*36*). Finally, over 6 billion people in the world have been infected with the protozoal pathogen *Toxoplasma gondii*, which can be fatal in infants and immunocompromised individuals (*37*). Thus the microbes and toxin used in H2P2 are important causes of human disease.

Beyond their role in human disease, the pathogens selected also exploit a wide range of host cellular processes to either kill the host cell or to create replicative niches within them. *C. trachomatis, S. enterica* serovar Typhi, serovar Typhimurium (wildtype and Δ*sifA* mutant, which escapes from the pathogen-containing vacuole at a greater rate into the host-cell cytosol (*38*)), and *T. gondii* are intracellular pathogens that employ diverse lifestyles. These microbes were engineered to express the green fluorescent protein (GFP) to allow quantitation of pathogen invasion, survival and replication, intercellular spread, and concurrent measurement of cell death by flow cytometry. B cells are known *in vivo* host cells for *Salmonella* invasion and replication (*39–41*) and the *C. trachomatis* strain that we utilize (L2) causes lymphogranuloma venereum (*42*), primarily an infection of the lymph nodes, where there is ample opportunity for interaction with B cells. Cell death was also measured as the readout for *Staphylococcus aureus* alpha toxin, a pore-forming toxin that causes cell lysis (*33*). Microbes were also tested for induction or suppression of cytokines (selected based on pilot studies of 41 cytokines) in infected LCLs. *S.* Typhimurium and *C. trachomatis-infected* cells were measured for 3 and 17 cytokines, respectively. The fungal pathogens *M. circinelloides* and *C. albicans* were included for their ability to induce FGF-2 in LCLs (*43*). Definitions for all 79 phenotypes as well as histograms for phenotypes are provided in the supplemental data (Table S1 and Figure S1). Importantly, 76 of 79 H2P2 phenotypes showed significant experimental repeatability based on measurements on LCLs from three different passages (Figure S2).

Hierarchical clustering of traits based on inter-individual variation confirmed the robustness of our measurements (Figure 1B). Levels of three cytokines (CXCL10 (IP-10), IL-10, and MDC) measured in uninfected cells with two different methods (ELISA at 24hrs and Luminex at 70hrs) showed strong correlation (R = 0.78 for CXCL10, R= 0.46 for IL-10, R=0.90 for MDC; Spearman correlation). The clustering of responses to *Salmonella* infection is consistent with previous findings: *S.* Typhimurium, Typhi, and Typhimurium Δ*sifA* cluster for the phenotype of invasion, as all utilize a similar type-3 secretion system for entry (*44*) (correlation to *S.* Typhimurium, R=0.69 for *S.* Typhi and R=0.90 for *S.* Typhimurium Δ*sifA)*. In contrast, the correlation is weaker for intracellular survival and replication phenotypes (correlation to *S.* Typhimurium replication from 3.5 hrs to 24 hrs, R=0.36 for *S.* Typhi and R=0.34 for *S.* Typhimurium Δ*sifA*), reflecting the different replications niches for Δ*sifA* (host cell cytoplasm) and wild-type Typhimurium (membrane bound vacuole) (*38*) or the use of a different repertoire of effectors by *S.* Typhi (*45*). In contrast, we observed almost no correlation between EBV copy number (from (*46*) for 284 LCLs also used in H2P2) and H2P2 traits (Figure S3; Table S2), indicating these phenotypes are not being driven by the LCL immortalization method. Thus, clustering based on phenotypic diversity verified the reliability of measurements and confirmed biological relatedness established by previous work from multiple groups.

### H2P2 traits are heritable based on parent-offspring regression and SNP-based heritability

If genetic differences regulate variation in cellular phenotypes, then the additive contributions of these differences to phenotypic variance can be estimated. We measured narrow-sense heritability (h^2^) with two independent and complementary methods: parent-offspring regression and SNP-based h^2^. Heritability based on parent-offspring regression is estimated as the slope of the regression line for offspring phenotypes vs. mid-parent phenotypes (*47*). We observed h^2^ estimates from −0.06 to 0.85 (average h^2^=0.33), with the largest h^2^ observed for *S.* Typhimurium-induced levels of IL-10 (h^2^=0.85±0.19, p=3.5×10^−15^) (Figure 1C; Figure S4; Table S3). The majority (64/79) of phenotypes showed significantly non-zero h^2^ by this method (p<0.05). In contrast, SNP-based h^2^, as implemented in the GCTA software package (*48*) with the Zaitlen modification for related individuals (*49*), calculates the proportion of variance that can be explained by all genotyped SNPs. With this method, we observed pedigree h^2^ ranging from 0.04 to 0.76 (average h^2^=0.36), and SNP-based h^2^ ranging from 0.02 to 0.66 (average h^2^=0.19) (Figure 1C; Table S3).

We observed high correlation between h^2^ estimated using the two different methods (R=0.58; p=2.9×10^−8^; Figure 1D). The strong correlation between the two estimates of h^2^, based on distinct statistical frameworks, provides additional evidence that LCLs provide a robust system to identify human SNPs that contribute to the heritability of cellular phenotypes.

### H2P2 reveals 17 genome-wide significant associations and enrichment for genic SNPs and regions of active chromatin

We performed a family-based GWAS in PLINK (*26, 27*) on 79 traits for 528 LCLs using dense genotyping information (15.5 million SNPs after imputation; see Methods). Across 79 traits, we observed 17 loci that reached a genome-wide significance threshold of p<5×10^−8^ (Figure 2A; Table 1; Figure S5). H2P2 demonstrated enrichment of associated SNPs for functional genome annotations. We used the GARFIELD package to calculate and visualize fold-enrichment of SNPs associated in H2P2 at variable p-value thresholds with different genomic features (*50*). In regards to SNP location, the greatest enrichment was observed for exonic SNPs (Figure 2B). The fold-enrichment was highest at the most stringent p-value threshold (p<1×10^−8^) for H2P2 traits (9.2-fold enrichment; p=0.10 by Fisher’s exact test) and was statistically significant when using a p<5×10^−8^ threshold for H2P2 traits (3.9 fold-enrichment; p=0.047). In contrast, there was depletion for intergenic SNPs at the p<5×10^−8^ threshold (0.66 fold-enrichment). H2P2-associated SNPs showed even greater enrichment for regions of active chromatin as annotated by DNase hypersensitivity peaks from the ENCODE project (*51*) (Figure 2C). Consistent with H2P2 being conducted in LCLs, the 2^nd^ greatest enrichment was observed for DNAase hypersensitivity peaks measured in the LCL GM06990 out of 424 cell types measured (at p<5×10^−8^, 5.7 fold-enrichment; p=3.8×10^−3^). Even stronger enrichment was noted at this threshold for human Th2 cells (7.3 fold-enrichment; p=3.5×10^−4^), and enrichment was observed for cells derived from most tissues (Figure 2C), consistent with LCLs being a relevant model for genetic analysis for multiple human cell types.

**Figure 2.**
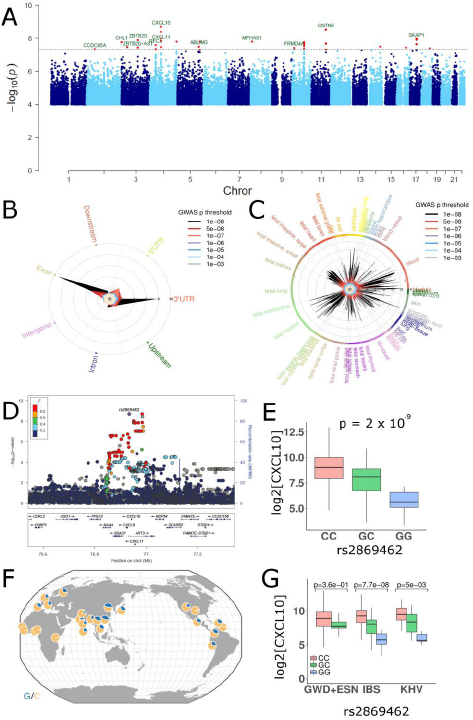
GWA of H2P2 reveals 17 genome-wide significant loci including a cis-cytokine-QTL near *CXCL10.* (A) A meta-Manhattan plot for 79 traits shows 17 peaks (red) with p<5×10^−8^ (dotted line). −log(p-values) were calculated using QFAM-parents in PLINK. (B) GARFIELD enrichment plot of SNP location demonstrates enrichment of SNPs associated with H2P2 phenotypes in exons, and 5’ and 3’ UTRs. SNPs associated with H2P2 traits at various p-value thresholds were plotted in the indicated colors and the height of the peak within each category indicates fold enrichment from 0 to 10. (C) GARFIELD enrichment plot of DNase hypersensitivity peaks demonstrates enrichment of SNPs associated with H2P2 phenotypes in active chromatin regions in multiple cell/tissue types. SNPs associated with H2P2 traits at various p-value thresholds were plotted in the indicated colors and the height of the peak within each category indicates fold enrichment from 0 to 14. (D) Regional plot around the *CXCL10* gene demonstrates association of rs2869462 with CXCL10 levels from *C. trachomatis-infected* cells. SNPs are plotted by position on chromosome 4 and −log(p-value) and color-coded by r^2^ value to rs2869462 from 1000 Genomes European data. (E) Genotypic medians, first and third quartiles (box), and maximum and minimum values (whiskers) for rs2869462 for CXCL10 levels from *C. trachomatis-infected* LCLs from all LCLs. (F) Map of rs2869462 allele frequencies (C = orange; G = blue) from Geography of Genetic Variants Browser (*81*). (G) Individual population genotypic median plots for rs2869462 for CXCL10 levels demonstrate C > G in all populations. P-values are from QFAM-parents in PLINK.

**Table 1.**
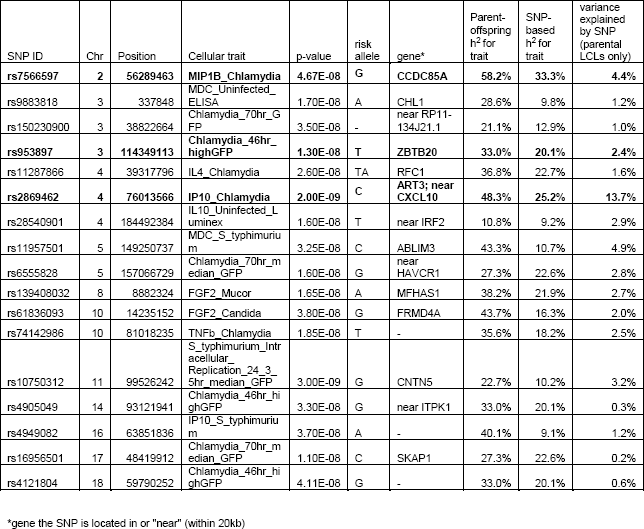
Genome-wide-significant H2P2 SNPs. A single SNP with the lowest p-value is listed for each peak. SNPs described in detail in the text are in bold.

| SNP ID | Chr | Position | Cellular trait | p-value | risk allele | gene* | Parent-offspring $h^2$ for trait | SNP-based $h^2$ for trait | variance explained by SNP (parental LCLs only) |
| --- | --- | --- | --- | --- | --- | --- | --- | --- | --- |
| **rs7566597** | **2** | **56289463** | **MIP1B_Chlamydia** | **4.67E-08** | **G** | **CCDC85A** | **58.2%** | **33.3%** | **4.4%** |
| rs9883818 | 3 | 337848 | MDC_Uninfected_ELISA | 1.70E-08 | A | CHL1 | 28.6% | 9.8% | 1.2% |
| rs150230900 | 3 | 38822664 | Chlamydia_70hr_GFP | 3.50E-08 | - | near RP11-134J21.1 | 21.1% | 12.9% | 1.0% |
| **rs953897** | **3** | **114349113** | **Chlamydia_46hr_highGFP** | **1.30E-08** | **T** | **ZBTB20** | **33.0%** | **20.1%** | **2.4%** |
| rs11287866 | 4 | 39317796 | IL4_Chlamydia | 2.60E-08 | TA | RFC1 | 36.8% | 22.7% | 1.6% |
| **rs2869462** | **4** | **76013566** | **IP10_Chlamydia** | **2.00E-09** | **C** | **ART3; near CXCL10** | **48.3%** | **25.2%** | **13.7%** |
| rs28540901 | 4 | 184492384 | IL10_Uninfected_Luminex | 1.60E-08 | T | near IRF2 | 10.8% | 9.2% | 2.9% |
| rs11957501 | 5 | 149250737 | MDC_S_typhimurium | 3.25E-08 | C | ABLIM3 | 43.3% | 10.7% | 4.9% |
| rs6555828 | 5 | 157066729 | Chlamydia_70hr_median_GFP | 1.60E-08 | G | near HAVCR1 | 27.3% | 22.6% | 2.8% |
| rs139408032 | 8 | 8882324 | FGF2_Mucor | 1.65E-08 | A | MFHAS1 | 38.2% | 21.9% | 2.7% |
| rs61836093 | 10 | 14235152 | FGF2_Candida | 3.80E-08 | G | FRMD4A | 43.7% | 16.3% | 2.0% |
| rs74142986 | 10 | 81018235 | TNFb_Chlamydia | 1.85E-08 | T | - | 35.6% | 18.2% | 2.5% |
| rs10750312 | 11 | 99526242 | S_typhimurium_Intracellular_Replication_24_3_5hr_median_GFP | 3.00E-09 | G | CNTN5 | 22.7% | 10.2% | 3.2% |
| rs4905049 | 14 | 93121941 | Chlamydia_46hr_highGFP | 3.30E-08 | G | near ITPK1 | 33.0% | 20.1% | 0.3% |
| rs4949082 | 16 | 63851836 | IP10_S_typhimurium | 3.70E-08 | A | - | 40.1% | 9.1% | 1.2% |
| rs16956501 | 17 | 48419912 | Chlamydia_70hr_median_GFP | 1.10E-08 | C | SKAP1 | 27.3% | 22.6% | 0.2% |
| rs4121804 | 18 | 59790252 | Chlamydia_46hr_highGFP | 4.11E-08 | G | - | 33.0% | 20.1% | 0.6% |

\*gene the SNP is located in or "near" (within 20kb)

### A large-effect cis-regulatory variant regulates CXCL10 levels

The strongest association in the H2P2 dataset was observed for rs2869462 with levels of the chemokine CXCL10 (also known as IP-10) following *Chlamydia* infection (p=2×10^−9^; Figure 2D, E). This SNP is located 7.5kb 3’ of the *CXCL10* coding sequence (Figure 2D) in a region that also encodes for two related chemokines, CXCL9 and CXCL11. These chemokines bind to CXC-chemokine receptor 3 (CXCR3), a G-protein coupled receptor that mediates inflammation by coordinating T-helper 1 (Th1) recruitment and activating effector cells during infection and autoimmunity (reviewed in (*52*)). Many cell types produce CXCL10 during infection, including activated B cells (*53–56*). Notably, the effect of this SNP is large: rs2869462 accounts for 13.7% of the variance in CXCL10 protein levels. While no dataset is available to replicate this association at the protein level, this SNP also demonstrated an association with *CXCL10* mRNA (p=7×10^−7^) in an independent set of 465 uninfected LCLs ((57); none of the LCLs overlap the H2P2 LCLs) (Figure S6).

Although rs2869462 was identified using genome-wide association of all 528 LCLs in our dataset, there were large differences in allele frequencies among the populations. The derived allele of rs2869462 (G) is present at the highest frequencies in Europe (28% in IBS) and Asia (28% in KHV) and is substantially lower in Africa (1.5% in ESN, 1% in GWD; Figure 2F). Strikingly, all populations demonstrated the same directionality of effect for the rs2869462 allele on CXCL10 levels (C > G) (Figure 2G).

### SNPs lead to pleiotropic effects on multiple pathogen-induced traits

Different pathogens can target common signaling pathways to establish an intracellular niche or to modulate immune responses. Therefore, we examined whether genome-wide significant hits were associated with just one trait or if they were associated with multiple traits, using either a threshold of p<0.05 or with Bonferroni multiple-test correction (p<6.3×10^−4^). All of the 17 genome-wide significant hits were associated with at least 4 H2P2 traits with the p<0.05 threshold (Figure 3A). However, several phenotypes are closely related and therefore these cross-phenotype associations do not reflect true pleiotropy, multiple unrelated effects due to the same gene (*58*). For example, rs2869462 had 5 cross-phenotype associations but 4 of these traits are based on CXCL10 levels (Figure 3B). Beyond the existence of pleiotropy, the pattern of which traits shared genetic associations provided additional insight (Figure 3C). A circle plot of cross-phenotype associations showed most cross-phenotype associations in H2P2 connect invasion, establishment of an intracellular niche, and intercellular spread. Traits that had high phenotypic correlation (Figure 1B) were more likely to have cross-phenotypic associations as expected (Figure 3D).

**Figure 3.**
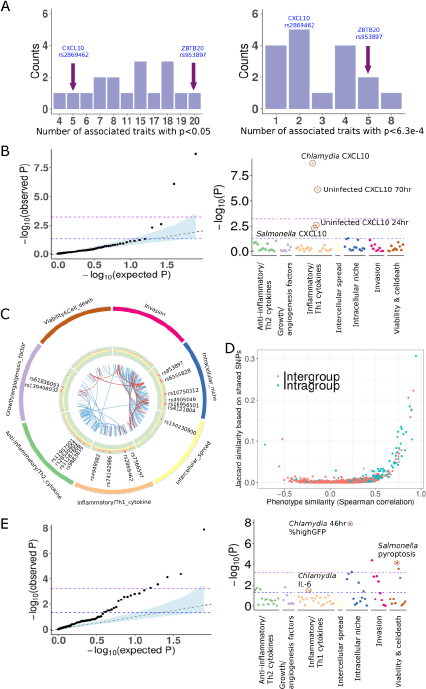
Cross-phenotype associations and pleiotropy are abundant among cellular host-pathogen traits. (A) Histograms of the number of cross-phenotype associations for the 17 H2P2 genome-wide significant hits. For the most strongly associated SNP in each GWAS peak, the number of traits with associations at p<0.05 and p<6.33×10^−4^ is shown. (B) QQ-plot and PheWAS plot for the association of rs2869462 with the 79 H2P2 phenotypes shows deviation from neutral expectation only for the 4 CXCL10 phenotypes. This is an example of cross¬phenotype associations but not pleiotropy. (C) Circle plot of 79 phenotypes by category and lines connecting traits that share the same genome-wide significant hit at p<1×10^−5^. (D) Plot of pairwise trait phenotypic similarity (Spearman correlation) vs. similarity of shared SNPs (Jaccard index). Traits that are more phenotypically similar have more shared SNPs with p< 1×10^−3^ for both traits. (E) QQ-plot and PheWAS plot for the association of rs953897 with the 79 H2P2 phenotypes shows deviation from neutral expectation for dozens of phenotypes, including traits in different biological categories in the PheWAS plot. This is an example of pleiotropy.

The greatest number of associated traits at p<0.05 occurred for rs953897, a SNP in the gene encoding the transcriptional repressor ZBTB20. Based on GTEx data, rs953897 is associated with *ZBTB20-AS1* transcript abundance (p=1×10^−9^) (*59*). While rs953897 is most strongly associated with high *C. trachomatis* burden at 46 hrs (p = 1.3×10^−8^), this SNP was associated with 20 H2P2 traits (p<0.05) and 5 traits using a multiple-test corrected threshold of p<6.3×10^−4^. A QQ plot comparing p-values for all traits for rs953897 confirmed the high degree of pleiotropy for this SNP, demonstrating strong deviation from neutrality towards lower p-values (Figure 3D). These associations even included other pathogens and biological processes, as the third most strongly associated trait was *S.* Typhimurium-induced pyroptosis (p=7.5×10^−5^; Figure 3D). Overall, out findings indicate that genetic variation in *ZBTB20* is linked to pleiotropic effects on multiple host-pathogen traits.

### Genetic variants impacting ZBTB20 expression affect the outcome of Chlamydia and Salmonella infections

The T allele of rs953897 was associated with both higher levels of *Chlamydia* replication (Figure 4A) and *Salmonella-induced* pyroptosis (Figure 4B). Reduction of *ZBTB20* expression by RNAi increased *Chlamydia* replication and pyroptosis, mimicking the effect of the T allele (Figure 4C, D). RNAi-mediated depletion of ZBTB20 also indicated that some phenotypes that did not reach statistically significant associations with rs953897 after multiple-test correction were nonetheless mediated by ZBT20. Specifically, expression of IL-6 after infection with *Chlamydia* (p=0.03), was *reduced* after *ZTB20* RNAi treatment, again mimicking the effect of the T allele (Figure 4E). The effect of ZBT20 depletion on IL-6 expression is not the result of decreased bacterial burden since *Chlamydia* replication was enhanced under these conditions. Thus, the association data and functional validation point to a role for ZBTB20 in the regulation of multiple infection-related phenotypes.

**Figure 4.**
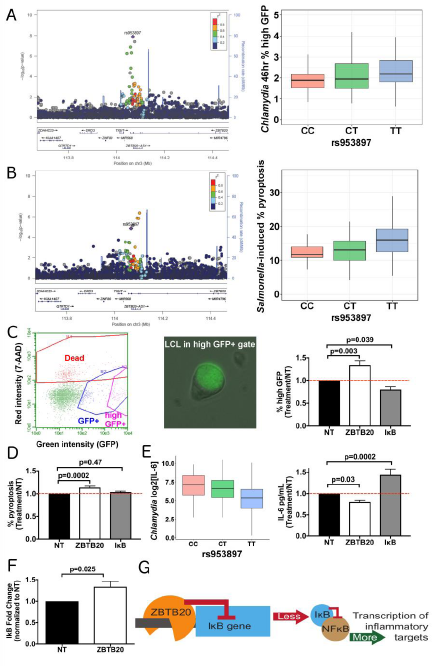
Genetic variation influencing *ZBTB20* regulates multiple host-pathogen traits. (A) Regional plot for *ZBTB20* demonstrates an association of rs953897 with *C. trachomatis* high GFP infected cells at 46 hrs (p=1.3×10^−8^). Genotypic medians plot of rs953897 with high GFP infected cells at 46 hrs in IBS LCLs. (B) Regional plot for ZBTB20 demonstrates an association of rs953897 with *S.* Typhimurium-induced pyroptosis (p=7.5×10^−5^). Genotypic medians plot of rs953897 with *S.* Typhimurium-induced pyroptosis in all LCLs. (C-F) LCL GM1761 was treated with non-targeting, ZBTB20 (53±9% knockdown), or IκB (83±2% knockdown) Accell RNAi for three days prior to infection. All cellular phenotypes measured were normalized to non-targeting values (treatment/non-targeting) prior to statistical analysis. (C) ZBTB20 and IκB regulate *C. trachomatis* replication. By 46hrs, *C. trachomatis* replication has resulted in a high GFP+ population of cells with an enlarged GFP+ *Chlamydia-containing* vacuole. ZBTB20 knockdown results in a greater percentage of high GFP+ cells, similar to what is seen with the T allele, while IKB knockdown produces fewer GFP+ enlarged vacuoles. Mean (±SEM) percentage of high GFP+ cells in non-targeting samples is 1.63% (±0.07%). (D) ZBTB20 regulates *Salmonella-induced* pyroptosis independent of IκB. ZBTB20 knockdown results in a greater percentage of pyroptotic cells, similar to what is seen with the T allele, while IKB knockdown shows no significant change in pyroptosis. Percentage of pyroptotic cells in non-targeting samples is 35.1% (±1.4%). (E) Both the T allele and ZBTB20 knockdown result in reduced IL-6. Genotypic median plot of rs953897 with *C. trachomatis*-induced IL-6 in all LCLs. ZBTB20 knockdown reduces IL-6 levels, while knockdown of IKB leads to increased levels measured at 70 hours. IL-6 levels from non-targeting LCLs were 186 pg/mL (±31.6 pg/mL). (F) ZBTB20 knockdown results in increased IκB mRNA. RNA was collected from nontargeting and ZBTB20 RNAi treated LCLs, cDNA was synthesized, and qPCR was conducted using TaqMan probes for 18s (control) and IκB (target). The ΔΔCT method was used to determine fold change of IκB. (G) Proposed model for ZBTB20 effect on proinflammatory targets. C-F were from 8-12 biological replicates from 2-4 experiments. p-values for C-E were generated from one-way ANOVA analysis while F was calculated by an unpaired parametric t-test.

ZBTB20 has been characterized as a transcriptional repressor during prenatal development in the liver and the brain (*60–62*). Rare protein-coding mutations in *ZBTB20* are responsible for Primrose syndrome, which has features as disparate as mental retardation, ossified external ears, and distal muscle wasting (*63*). We hypothesized that one ZBTB20 target gene could regulate a pathway that impacted multiple biological processes related to pathogen immunity. An attractive target, I*κ*B *(NFKBIA)*, the canonical suppressor of NF*κ*B signaling, is subject to ZBTB20 transcriptional repression (*64*). Reduction of ZBTB20 expression in LCLs by RNAi causes increased expression of I*κ*B (Figure 4F). An increase of I*κ*B should cause inhibition of NF*κ*B signaling, resulting in increased *Chlamydia* replication but decreased expression of pro-inflammatory cytokines including IL-6. Consistent with this prediction, depletion of I*κ*B by RNAi decreased *Chlamydia* replication and increased IL-6 production (Figure 4C, E). In contrast, depletion of I*κ*B did not impact *Salmonella-induced* pyroptosis, indicating an I*κ*B-independent mechanism (Figure 4D). These observations point to multiple roles for ZBTB20 in regulating cellular functions during infection, both through suppression of regulators of signaling pathways, such as NF*κ*B (Figure 4J), but also through regulation of other unidentified targets.

### SNPs linked to CXCL10 expression are also associated with inflammatory bowel disease

We determined if SNPs associated with cellular traits in H2P2 were associated with human disease. Consistent with the rs2869462 C allele being associated with increased CXCL10 and inflammation, we discovered that this allele is a previously unrecognized inflammatory bowel disease (IBD) risk allele. CXCL10 inhibitory antibodies have undergone phase II clinical trials for both sub-types of IBD, Crohn’s disease (CD) and ulcerative colitis (UC) (*65, 66*), based on evidence from animal models (*67–69*) and of elevated CXCL10 in patients with CD (*70*) and UC (*71*). Comparison to GWAS summary statistics from the IBD Genetics Consortium meta¬analysis of 12882 IBD cases and 21770 controls (*72*) demonstrated that rs2869462 is associated with IBD (p=1.7×10^−4^; OR=1.08), as well as with CD (p=1.9×10^−3^; OR=1.09) and UC subtypes (p=0.016; OR=1.06). The cases in the subtype analysis are exclusive of one another and therefore demonstrate an association of rs2869462 with two different cohorts of IBD cases (though most controls are shared). Furthermore, the direction of association is consistent with high levels of CXCL10 (the C allele) being associated with greater risk of IBD.

The availability of GWAS summary statistics from this dataset allowed us to conduct a formal colocalization analysis to determine whether the CXCL10 protein level signal was the same as the IBD signal. We utilized the COLOC package which uses a Bayesian framework to determine whether GWAS signals in the same region are likely due to the same causal variant (*73*). The posterior probability that both CXCL10 protein level and IBD share the same causal variant is as high as 0.80 for the region, with rs2869462 being singled out as the SNP with the greatest posterior probability as being causal (Table S4). Comparison of regional plots of association along with the colocalization analysis indicate that the same LD block is associated with both CXCL10 levels and risk of IBD (Figure 5A).

**Figure 5.**
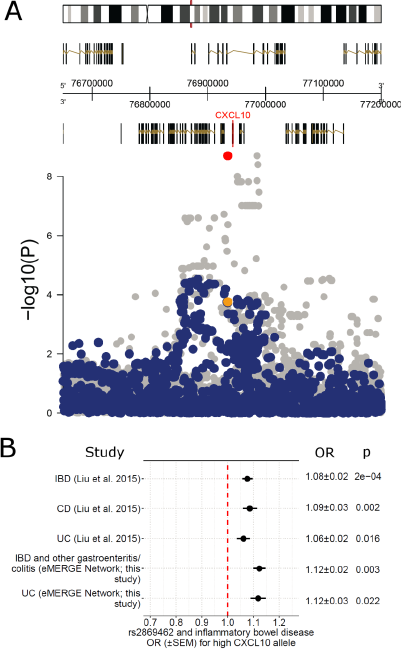
SNPs associated with high CXCL10 in H2P2 are also associated with increased risk of IBD. (A) Overlaid association plots of the CXCL10 region on chromosome 4 demonstrates colocalization of signals for *Chlamydia-infected* CXCL10 levels from H2P2 (grey) and IBD GWAS (blue) (*72*). rs2869462 from the two studies is highlighted in red and yellow. Colocalization analysis indicates an 80% probability that the peaks for CXCL10 level and IBD are due to the same causal variant (Table S4). (B) OR plot for rs2869462 and IBD, CD and UC based on data generated in (*72*) and replication of the association with “IBD and other gastroenteritis and colitis” and UC from the eMERGE Network. The high CXCL10 allele (C) is associated with increased odds of IBD.

We independently tested for this association using electronic medical record (EMR) data from the eMERGE Network (*74*). The eMERGE dataset holds genotype-phenotype correlations of >80,000 individuals with phenotypes assigned based on ICD-9 patient billing codes (*25*). We tested for association with the codes for “inflammatory bowel disease and other gastroenteritis and colitis” and the more restrictive code for “ulcerative colitis.” rs2869462 was associated with both phenotypes in the predicted direction (p=0.003; OR=1.12, C allele for IBD, p=0.02; OR=1.12, C allele for UC) (Figure 5B). Therefore by first identifying a SNP as associated with CXCL10 levels following an LCL model of *C. trachomatis* infection, we have discovered and replicated a new IBD risk allele.

### H2P2 SNPs are associated with disease in PheWAS of electronic medical record traits

Next, we systematically generated hypotheses regarding the effects of the H2P2 genome-wide significant hits on human disease by employing a PheWAS (phenome-wide association study) approach, looking for associations across a large set (1338) of clinical measurements and diseases cataloged in eMERGE (*25*). We found 5 of 16 H2P2 genome-wide significant hits surpassed the multiple test corrected significance threshold with at least 1 EMR phenotype (Figure 6A; Table S5; 1 H2P2 SNP was not in the eMERGE dataset and had no good proxy based on LD).

**Figure 6.**
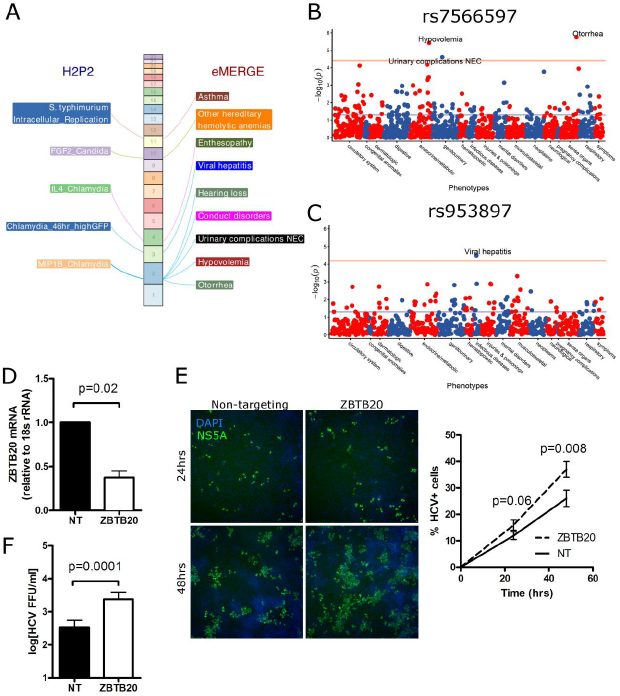
PheWAS of H2P2 hits reveals connections to human disease including an association of *ZBTB20* and viral hepatitis. (A) Chromosome landscape of H2P2 genome-wide significant hits that demonstrate significant associations with eMERGE phenotypes, after Bonferroni correction for the number of eMERGE phenotypes. The five SNPs demonstrating significant associations are described in Table S5. (B) (C) eMERGE PheWAS plots of rs7566597 and rs953897. eMERGE phenotypes are grouped into functional categories. Y-axis is −log(p-value), with the red line indicating Bonferroni corrected p-value = 0.05. (D) Huh7 cells were transfected with non-targeting or ZBTB20 siRNA for two days prior to infection. qPCR demonstrated significant knockdown of ZBTB20. (E) ZBTB20 suppresses HCV infection. The percentage of cells infected with HCV was assessed at 24 and 48 hpi by immunofluorescence for HCV protein NS5A (green) and staining with DAPI (blue) for total cells, with automated quantification by Cellomics. ZBTB20 depletion causes the percentage of HCV-infected cells to increase more rapidly over time. (F) ZBTB20 suppresses HCV production. Supernatants collected at 72 hpi were used in a HCV focus forming assay to determine the concentration of productive HCV particles. ZBTB20 depletion causes a 7-fold increase in HCV focus forming units (FFUs/ml). For D-F, mean (±SEM) are shown and p-values are from a paired t-tests from three separate experiments.

The SNP that had the greatest number of PheWAS associations was rs7566597. This SNP, associated with the H2P2 phenotype of *Chlamydia-infected* levels of the chemokine MIP-ip (macrophage inflammatory protein-1beta; CCL4), was associated with five clinical traits. The most significant association (p =1.76×10^−6^; p=0.0024 after Bonferroni; OR=1.40) was observed with otorrhea (Figure 6B). Otorrhea is ear drainage most commonly caused by an ear infection. Notably, MIP-1β is elevated in fluid from patients with middle ear infections (*75*), mice with genetic predisposition to middle ear infections (*76*), and primary middle ear epithelial cultures infected *in vitro* with either influenza or *Streptococcus pneumoniae* (*77*). The directionality of this association (G allele associated with both higher MIP-1β levels and increased risk of otorrhea), and the fact that genotype is fixed prior to disease, lead to the hypothesis that the rs7566597 G allele causes higher levels of MIP-1β to increase risk of otorrhea.

Similarly, rs953897 in *ZBTB20* was associated with viral hepatitis (Figure 6C; p=3.17×10^−5^; p=0.025 after Bonferroni; OR=1.20). Most of the 1654 viral hepatitis cases in this cohort are due to hepatitis C virus (HCV). To further test this association experimentally, we performed RNAi against *ZBTB20* in Huh7 human hepatocytes. Depletion of *ZBTB20* mRNA (Figure 6D) increased the percentage of HCV infected cells over time (Figure 6E) and increased the amount of infectious virus by 7-fold (Figure 6F). Future mechanistic and clinical studies will be required to further validate the association and determine how genetic variation in *ZBTB20* affects risk of viral hepatitis. However, this example demonstrates that coupling H2P2 with PheWAS of clinical traits can lead to hypotheses that can be quickly tested in the most clinically relevant cell type for that particular disease.

## Discussion

With H2P2, we have coupled the ability of pathogens to influence genetic diversity in human populations to their use as cellular probes to elucidate mechanisms of disease. This cellular GWAS approach provides: 1) biomolecules and proteins that could serve as possible biomarkers and therapeutic targets and 2) a cellular model to validate and dissect mechanisms of how the genetic variant regulates the cell biological process now connected to disease. We have developed an H2P2 database and web interface to allow for exploration of this rich dataset by the research community (http://h2p2.oit.duke.edu).

Cellular GWAS studies have also been performed on inter-individual variation in levels of immune cell subtypes, cell surface protein expression, and cytokine levels (*78–80*). However, Hi-HOST is unique in using live pathogens to induce complex cellular phenotypes, such as cell death and invasion, providing surrogate phenotypes that are intermediate between molecular phenotypes of gene/protein expression and human population studies of disease. Additionally, H2P2 utilizes cells derived from multiple human populations. Notably, allele frequencies for rs2869462 and rs953897 vary greatly across the globe. These differences may play an important role in susceptibility of different populations in cellular responses and disease. Finally, our use of parent-offspring trios allowed us to make estimates of h^2^ through both parent-offspring regression and SNP-based methods.

Our h^2^ estimates are consistent with a large fraction of variation in cellular responses as being genetically determined. For parent-offspring regression 64 of 79 traits had significantly non-zero h^2^ (p<0.05). While relatively small sample size led to large standard errors for SNP-based estimates of h^2^, the two estimates correlated quite strongly (r=0.58; p=2.9×10^−8^). Our estimates of h^2^ are similar to other reports for immune-related traits. Orru *et al*. examined levels of 95 immune cell types and found a mean h^2^ of 41% (range 3-87%) (*79*). Our estimates of h^2^ for cytokine levels as well as for more complex host-pathogen phenotypes such as *Chlamydia* intracellular replication (46hr % high GFP; 33% by parent-offspring regression and 20% by SNP-based heritability) are consistent with a strong genetic basic for variation in immune cell traits and host-pathogen interactions, though environment also has a large effect. Furthermore, our ability to identify SNPs strongly associated with these traits that can explain a sizable portion of the h^2^ further confirms the importance of common SNPs in contributing to the h^2^ of molecular and cellular pathogen-induced traits.

Integrating H2P2 with human genetic association data from published studies and the eMERGE Network revealed how genetic variants impacting cellular traits also influenced human disease. For rs2869462, we found that this SNP associated with CXCL10 levels is also a previously unrecognized risk factor for IBD. While over a hundred IBD risk alleles have been identified (*72*), the fact rs2869462 was associated with levels of CXCL10 in H2P2 may make genotyping of this SNP clinically actionable if coupled to anti-CXCL10 therapy. While anti-CXCL10 demonstrated some benefit in phase 2 trials, neither study met statistical significance for its primary endpoint (*65, 66*). We hypothesize that rs2869462 genotype might be a predictive biomarker for identifying the genetic subtype of patients who show the greatest benefit. This example, spanning molecular phenotype, human disease, and clinical utility serves as a template for how we envision the H2P2 web portal being used to make similar discoveries.

For *CXCL10, ZBTB20,* and other genes implicated by H2P2, there are numerous associations that do not reach genome-wide significance but are undoubtedly true-positives based on highly related phenotypes or experimental evidence. Indeed, our lab has previously pursued non-genome-wide significant hits revealed by Hi-HOST, resulting in a new metabolite biomarker for sepsis (*23*) and a new potential therapeutic strategy for typhoid fever (*24*). However, to fully illuminate how genetic variation contributes to the pathophysiology of disease will require the engagement of the research community with the H2P2 web portal and other similar datasets. These users, already experts on particular genes and/or cellular pathways, would be well-equipped to then validate and discover the mechanisms underlying these associations. Thus, H2P2 provides a hypothesis-generating engine for identifying new biomarkers and therapeutic strategies. Future studies will expand the panel of stimuli and the cell types used to create a more complete picture of how cellular traits impact human health and disease, an important step towards a future of more personalized care.

## Author Contributions

All authors critically reviewed the manuscript and contributed input to the final submission. DCK, LW, KJP, ALA, IBS, GDW, TB, SMH, RHV, and DRC wrote the manuscript. DCK, LW, KJP, ALA, JH, SCL, SMH, RHV, and DRC contributed to strategy and project planning. KJP, DCK, JRB, RES, GDW, AMG, KDG, and SCL carried out experiments and analysis. LW, IBS, RJC, tEM, GPJ, JCD, DRC, and DCK planned and carried out computational analysis and provided datasets. TB, AI, LW, MRD, and DCK designed and implemented the H2P2 database and web portal.

## Acknowledgments

Funding: LW, KJP, RES, KDG, ALA, YC, and DCK were supported by NIH R01AI118903. LW, KJP, JRB, RES, AMG, RHV, and DCK were supported by the Ecopathogenomics of Sexually Transmitted Infections (EPSTI) Cooperative Research Center (NIH U19AI084044). DCK was also supported by a Duke University Whitehead Scholarship, the Butler Pioneer Award, and a Duke MGM Pilot Award. GDW and SMH were supported by NIH R01AI125416 and R21AI124100 and the Burroughs Wellcome Fund. IBS, RJC, JCD, and DRC were supported by eMERGE Network (Phase III). This phase of the eMERGE Network was initiated and funded by the NHGRI through the following grants: U01HG8657 (Group Health Cooperative/University of Washington); U01HG8685 (Brigham and Women’s Hospital); U01HG8672 (Vanderbilt University Medical Center); U01HG8666 (Cincinnati Children’s Hospital Medical Center); U01HG6379 (Mayo Clinic); U01HG8679 (Geisinger Clinic); U01HG8680 (Columbia University Health Sciences); U01HG8684 (Children’s Hospital of Philadelphia); U01HG8673 (Northwestern University); U01HG8701 (Vanderbilt University Medical Center serving as the Coordinating Center); U01HG8676 (Partners Healthcare/Broad Institute); and U01HG8664 (Baylor College of Medicine). U01HG004438 (CIDR) and U01HG004424 (the Broad Institute) served as eMERGE Genotyping Centers. SCL was supported by NIH/NIAID R03 AI11917 and UTSA Research funds. JH was supported by NIH/NIAID R37 MERIT Award AI39115-20, NIH/NIAID R01 AI50113-13, and an award from Astellas. TB, AI, and MRD were supported by Duke Research Computing. We thank Dr. Yanlu Cao for performing FGF-2 ELISAs. We thank Luke C. Glover for help in plotting association data. We thank Dr. Gregory D. Sempowski and the Duke Immune Reconstitution & Biomarker Analysis Shared Resource in performing pilot Luminex measurements. We thank Dr. Charles Rice for the HCV NS5A antibody and the Duke Functional Genomics Shared Resource for use of the Cellomics ArrayScan. We thank Drs. Shelton Bradrick, David Tobin, Ana-Maria Xet-Mull, Andrew Alspaugh, and Jennifer L. Trenor for pilot studies in characterizing infection of various pathogens into LCLs. DNA image from Figure 1 is from the National Human Genome Research Institute Image Gallery. The content of this manuscript is solely the responsibility of the authors and does not necessarily represent the official views of the National Institutes of Health or other funding sources.

## Methods

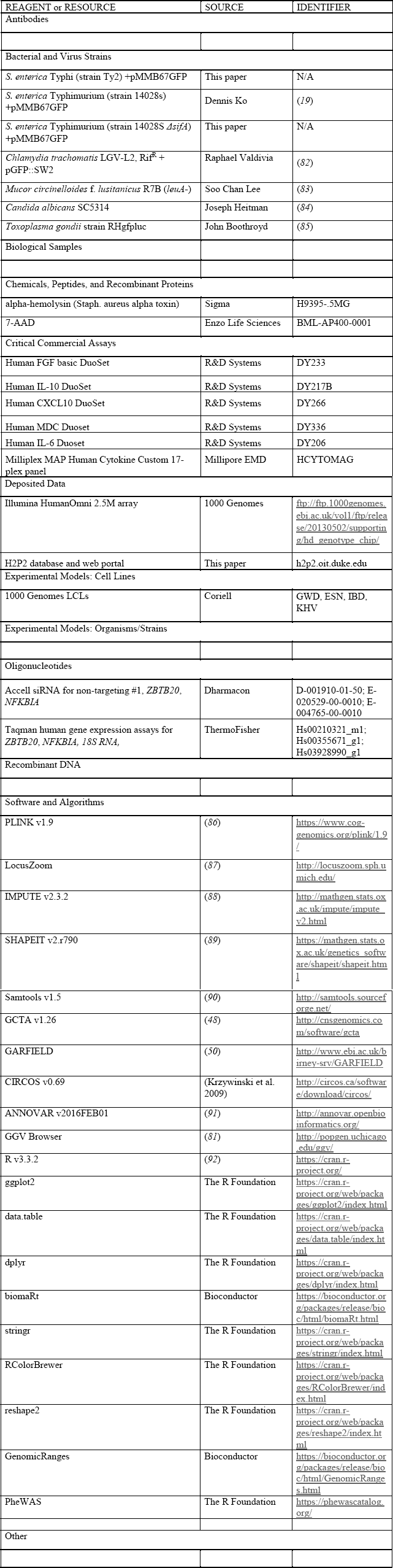
KEY RESOURCES TABLE.

## CONTACT FOR REAGENT AND RESOURCE SHARING

Further information and requests for reagents should be directed to and will be fulfilled by the Lead Contact, Dennis Ko.

## EXPERIMENTAL MODEL AND SUBJECT DETAILS

### Cells

1000 Genomes LCLs (528; all trios) from ESN (Esan in Nigeria), GWD (Gambians in Western Divisions in the Gambia), IBS (Iberian Population in Spain), and KHV (Kinh in Ho Chi Minh City, Vietnam) populations were purchased from the Coriell Institute. LCLs were maintained at 37°C in a 5% CO_2_ atmosphere and were grown in RPMI 1640 media (Invitrogen) supplemented with 10% fetal bovine serum (FBS), 2 mM glutamine, 100 U/ml penicillin-G, and 100 mg/ml streptomycin.

Human hepatoma Huh7 cells were grown in DMEM (Mediatech) supplemented with 10% fetal bovine serum (HyClone), 2.5 mM HEPES, and 1× non-essential amino acids (complete, cDMEM; Thermo Fisher Scientific). The identity of the Huh7 cell line was verified using the Promega GenePrint STR kit (DNA Analysis Facility, Duke University), and cells were verified as mycoplasma free by the LookOut Mycoplasma PCR detection kit (Sigma). Infectious stocks of a cell culture-adapted strain of genotype 2A JFH1 HCV were generated and titrated by focus-forming assay (FFA), as described (*93*).

## METHOD DETAILS LCL

### screening

LCLs were received from Coriell and cultured for 8 days prior to assays. LCLs were counted with a Guava Easycyte Plus flow cytometer (Millipore). LCLs were washed once with RPMI 1% FBS and then plated out in RPMI 10% FBS at 200,000 cells/200μl for Salmonellae, 100,000 cells/100μl for fungi, and 40,000 cells/100μl for *S. aureus* alpha toxin, *C. trachomatis*, and *T. gondii*. Cells were passaged at 150,000/ml in 20ml total volume for three days.

### *C. trachomatis* infection

*C. trachomatis* LGV-L2 Rif^R^-pGFP::SW2 was grown and purified as previously described *(94). C. trachomatis* was added at MOI 5 in 100μl assay media, mixed by multichannel pipetting, and centrifuged onto cells at 3000 RPM for 30min at 4°C. At 27, 46, and 70hrs, cells were mixed and 25μl was taken for flow cytometry measurement (4000 cells). 25μl of supernatant at 70hrs was measured by Luminex assay for 17 cytokines.

### *Salmonella* infection

Salmonellae were tagged with an inducible GFP plasmid [pMMB67GFP from (*95*)]. *sifA* deletion mutants was constructed with lambda red (*96*) and verified by PCR. Assaying LCLs for Salmonellae infection was conducted as previously described (*19*). Overnight bacterial cultures were subcultured with a 1:33 dilution and grown for 2 hr 40 min at 37°C. Invasion was conducted for 1hr at a multiplicity of infection (MOI) of 10 for *S.* Typhi and MOI 30, followed by addition of gentamicin (50μg/ml) for 1hr, and then culture was split into two separate cultures of 60μl of cells with 140μl of media to dilute gentamicin (15μg/ml) and allow for collection at two timepoints. IPTG (1.4mM) was added to turn on GFP expression for 75 min prior to 3.5hr and 24hr timepoints. For the 3.5hr timepoint, 150μl of cells were stained with 7-AAD (7-aminoactinomycin D; Enzo Life Sciences) and green and red fluorescence of 7000 cells was measured on a Guava Easycyte Plus flow cytometer (Millipore). For the 24hr timepoint, cells were spun down and 2 aliquots of 55ul of supernatant was removed and stored at -80C for subsequence IL10 (25μl), CXCL10 (25μl), and MDC (4μl) ELISAs (R&D Systems). 55μl of cells were stained with 7-AAD and measured by flow cytometry.

### Fungal infection

The *Mucor circinelloides* f. *lusitanicus* R7B *(leuA*-) (*83*) strain and *Candida albicans* SC5314 (*84*) strain were used as wild-type to test the expression of FGF-2 from the LCLs. Leucine autotropism (*leuA^-^*) was found not to impact virulence (*97*). To prepare *Mucor* spores, potato dextrose agar (PDA, 4 g potato starch, 20g dextrose, and 15 g agar per liter) was inoculated and incubated for 4 days at 26°C in the light. To collect spores, sterile deionized distilled water was added onto the plates and spores were released by gently scraping the colonies with a cell spreader. Spores were counted by using a hemocytometer. To prepare *C. albicans* yeast, yeast dextrose broth (10 g yeast extract, 20 g peptone, 20 g glucose per liter) was inoculated and incubated at 30°C by shaking at 250 rpm overnight. The yeast cells were quantified by using a hemocytometer. To co-culture with LCLs, all fungal cells were washed with sterile PBS twice. Fungi were added at MOI 1 in 10μ1 and incubated for 24 hrs. Culture supernatant was collected and stored at -80°C for later FGF-2 ELISA analysis (R&D Systems).

### *S. aureus* toxin treatment

LCLs were treated with alpha-hemolysin (Sigma) at 1μg/ml for 23hrs. Cells were mixed and cell death quantified by 7-AAD staining (concentration) and flow cytometry.

### *T. gondii* infection

*T. gondii* strain RHgfpluc was grown on confluent human foreskin fibroblast cells. The infected cells were then scraped and transferred to a 50ml polystyrene tube and centrifuged at 500 × g for 10 minutes at 4°C. Pellet was resuspended in 3 ml of PBS and the suspension was aspirated three times using a 20g needle attached to a 10ml syringe. 30 ml of PBS was added and centrifuged at 500 × g for 10 min at 4°C. Supernatant was removed, and pellet resuspended in 5ml of PBS. Concentration of a 1:200 dilution was determined by flow cytometry and added at MOI 2 to cells. At 5, 30, and 48 hrs infection, cells were mixed, and 25μl taken for measuring 4000 cells by flow cytometry.

### LCL RNAi experiments

LCLs (2×10^5^ cells) were treated for three days in 500μl of Accell media (Dharmacon) with either non-targeting Accell siRNA #1 or an Accell SmartPool directed against human *ZBTB20* or *NFKBIA* (1μM total siRNA; Dharmacon). Prior to infection, cells were plated at 1×10^5^ in 100μl RPMI complete media (without antibiotics) in 96-well plates. Infections were conducted as described above.

### HCV infection experiments

Huh7 cells were seeded in 12-well plates at a density of 1×10^5^ cells per-well in cDMEM and transfected the next day using 9μL RNAiMAX (Thermo) and 3μL of the indicated 10μM siRNA (siGenome Smartpool (Dharmacon)), in Optimem (Thermo). Four hours post-transfection, the transfection mixture was removed and 1ml fresh cDMEM was added. HCV infections were performed at an MOI of 0.3 for 24, 48, or 72 hr. For each condition, duplicate wells were either infected or mock-infected, RNA was harvested from one well and the other utilized for visualization of infected cells. Supernatant was collected from both wells for virus titration.

HCV focus forming assay. Serial dilutions of supernatants collected from non-targeting or ZBTB20-targeting siRNA treated cells collected 72hpi were used to infect naive Huh7.5 cells in triplicate wells of a 48-well plate. At 48 hpi, cells were fixed, permeabilized, and immunostained with HCV NS5A antibody (1:500; gift of Charles Rice, Rockefeller University). Following binding of horseradish peroxidase (HRP)-conjugated secondary antibody (1:500; Jackson ImmunoResearch), infected foci were visualized with the VIP Peroxidase Substrate Kit (Vector Laboratories) and counted at 40× magnification.

Visualization of HCV infected cells. 48 hours post-transfection, Huh7 cells treated with non-targeting or ZBTB20-targeting siRNA were infected with HCV (MOI 0.3) or mock-infected. 24 or 48 hours post-infection, cells were fixed in 4% paraformaldehyde in PBS, permeabilized with 0.2% Triton X-100 in PBS, and blocked with 3% BSA in PBS, then immunostained with HCV NS5A antibody (1:1,000), washed 3× in PBS-Tween, then visualized with AlexaFluor-488 Donkey anti-Mouse secondary antibody (1:1000, Thermo). Cell nuclei were stained with DAPI in the first of three PBS-Tween washes following the addition of secondary antibody. Two-color images were collected with the Cellomics ArrayScan VTI HCS (Thermo), at 20x magnification in the Duke Functional Genomics Shared Resource. 10 fields of per-well, per-condition were acquired and the percentage of identified nuclei with detectable NS5A staining was quantified using VHSview software (Thermo).

## QUANTIFICATION AND STATISTICAL ANALYSIS

### Phenotype repeatability

Repeatability of each cellular trait was calculated to measure the variation among 3 independent experiments. The inter-and within-individual component of variance was calculated by fitting to one-way ANOVA. The estimated within-individual component of variance gave the repeatability coefficient.

### Testing effect of EBV copy number on H2P2 cellular phenotypes

EBV relative copy numbers were retrieved for 1753 LCLs (*46*), of which 284 cell lines overlapped with H2P2 samples (73 ESN; 105 GWD; 106 IBS). Prior to analysis, H2P2 cell phenotypes were averaged from the three experiment replicates and then transformed to Z-scores within each experimental batch. EBV loads were standardized to Z-score within each population. Correlation between H2P2 cellular traits and EBV loads were tested using linear regression with population as a covariate.

### Genotype and imputation

Genotypes for 1000 Genome LCLs (*16*) were from Illumina HumanOmni 2.5M array (905,788 SNPs; see details in STAR resource table). Genome-wide imputation of autosomal genotypes with 1000 genome Phase 3 haplotype as reference panel were performed through two steps, a pre-phasing step using SHAPEIT2 (*89*) and an imputation step using IMPUTE2 (*98*). Imputed genotype was further filtered by imputation accuracy score (IMPUTE’s INFO) < 0.9 and minor allele frequency < 0.01. A total of 339 samples overlap with 1000 genome Phase 3 individuals. We merged those direct sequenced genotypes from 1000 genome Phase 3 project into our imputed genotypes. We eventually obtained 15,581,278 SNPs (8,817,925 SNPs have minor allele frequency ≥ 0.05). The human genome reference assemble (GRCH37/hg19) was used for all analysis.

### Phenotype-and SNP-based heritability analysis

Two different methods were applied to estimate heritability. The parent-offspring (PO) regression method estimated additive heritability exclusively using phenotypic values. Linear regression of child against average of parents was performed, and the slope was used as a heritability estimator. Batch was incorporated as a covariate. Genotype-based heritability was estimated using the GCTA GREML method (Yang et al. 2012). Autosomal SNPs with minor allele frequency filtering of 0.05 were used to create a genetic relationship matrix (GRM). Zaitlen and colleagues developed a method, “big K/small K”, which enables to precise estimate heritability by jointly using closely related and unrelated individuals (*49*). Following Zaitlen’s method, variance explained by genome-wide SNPs (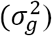) was then estimated for each cellular trait. The Zaitlen modification provides joint estimates of 1) h^2^ based on pedigree relatedness and 2) h^2^ based on inferred relatedness from genome-wide SNPs.

While the standard error for SNP-based h^2^ estimates were quite large, we nonetheless observed very strong correlation between these estimates and the parent-offspring h^2^ estimates (see Figure 1D). We estimated h^2^ based on the analysis of LCLs from all populations in H2P2 to increase the precision of our estimates by including more individuals. Although h^2^ is a population-specific parameter, h^2^ are often quite similar across different populations and even species (*99*). Nonetheless, we include estimates and standard errors for the combined analysis as well as individual populations in Table S2.

### Genome-wide association analysis

Genome-wide association analysis was conducted with PLINK v1.9 (*26*). Analysis was carried out using the QFAM-parents approach with adaptive permutation and a maximum of 10^9^ permutations. The QFAM procedures implemented in PLINK use linear regression to test for association while employing permutation of within-and between-family components separately to control for family structure (*27*).

### Enrichment analysis

Enrichment analyses were carried out using GARFIELD (*50*). GARFIELD has predefined a total of 1005 features from ENCODE and the NIH Roadmap project, and applies generalized linear regression models while accounting for the effects of linkage disequilibrium (LD), minor allele frequency, and local gene density. The GWAS summary statistics were used to quantify fold-enrichment against predefined annotation features at different GWAS p-value thresholds.

### PheWAS Analysis

Testing for association of H2P2 genome-wide hits with clinical phenotypes was performed with the eMERGE biobank dataset of 83,717 individuals from 12 contributing medical centers (*1–4*) with ICD-9 derived PheWAS codes (*5, 6*). A merged set of unified variant genotypes across 78 batches of samples with different genotype platforms (e.g. various Illumina and Affymetrix arrays) was produced by imputation using the Michigan Imputation Server (MIS) with the HRC1.1 haplotype reference set (7-9). Sixteen variants were selected for PheWAS based on association in H2P2 and their inclusion in the imputed eMERGE biobank dataset. The PheWAS codes were defined by query of the ICD-9 electronic medical record datasets of the contributing medical centers. Two types of PheWAS code phenotypes were used in the association to ascertain more chronic versus singleton diagnoses: the minimum code count of one (mcc1) ICD-9 code to define an individual as a PheWAS code case, and minimum code count of two (mcc2) instances to define an individual as a chronically represented case. In the mcc2 cases, individuals were excluded from analysis if they only had one instance of the ICD-9 code. If there were less than 500 cases we did not include the ICD-9 derived PheWAS code in analysis because it would likely be underpowered and impact the multiple testing correction. We also did not include medical centers that had low ascertainment of the ICD-9 by excluding medical centers which had less than 10 cases. This resulted in 1,338 for mcc1 and 788 for mcc2 phenotypes being included in the analysis. We used the PLINK1.9 identity by descent genome file to find the set of unrelated individuals to bring forward for analysis. PheWAS association was implemented in the R *glm()* logistic regression of the case-control data and plotting was carried out using the PheWAS R package (*5*). The covariates of gender and the PLINK1.9 computed 1 and 2 principal components from the pruned (>5% minor allele frequency, genotype and sample missingness > 0.1 and LD r-square<0.7.) genome wide imputation variant genotypes were included in the regressions. The p.adjust() R function with Bonferroni methods was used to adjust p-values of the tested PheWAS codes within a particular SNVs sets of tests for multiple comparisons. Bonferroni of less than 0.05 was used as a significance threshold.

### Colocalization analysis

Colocalization analysis was performed using R “coloc” v2.3.1 package (available at http://cran.r-project.org/web/packages/coloc). This software applies a Bayesian framework to estimate the posterior probability of genomic variants affecting both cellular trait and disease based on pre-computed GWAS p values, odds ratios, and minor allele frequencies. We ran colocalization on a 400-kb region centered on the focus SNP rs2869462 using default COLOC parameters (P1=P2=1×10^−4^; P12=1×10^−5^). Summary statistics of IBD GWAS (*72*) were obtained from www.ibdgenetics.org.

### Gene expression analysis on 1000 genome RNAseq project

Gene expression data of 465 individuals (*57*) were obtained from the EBI website (https://www.ebi.ac.uk/Tools/geuvadis-das/). The rs2869462 genotype data was downloaded from the 1000 genome project (*16*). Effects of rs2869462 on *CXCL10* gene expression were tested by linear regression on both combined populations and individual population.

### Descriptive statistics and visualization

Descriptive statistics were performed with GraphPad Prism 6 (GraphPad Software, US) and with R (*92*). QQ plots were plotted using quantile-quantile function in R. Regional Manhattan plot were made using LocusZoom (*87*). Circos v0.69 was used to visualize the shared SNPs among different groups. The size of each study or number of replicates, along with the statistical tests performed can be found in Figure Legends. All numerical data are presented as the mean ± SEM (standard error of mean).

## DATA AND SOFTWARE AVAILABILITY

All H2P2 data is available for browsing and download at http://h2p2.oit.duke.edu The H2P2 application server is running Red Hat Enterprise Linux Server v7.4, Apache v2.4.6, Shiny Server v1.5.3.838, R v3.4.1, and Microsoft ODBC Driver 13 for SQL Server. Implemented R packages include shiny v1.0.3, RODBC v1.3-15, ggplot2 v2.2.1, d3heatmap v0.6.11, and DT v0.2. The H2P2 database server is running MS Windows Server 2016 and MS SQL Server 2016. Large volume tables and indexes (2.5 billion GWAS observations and 8.5 billion genotype observations) were partitioned for improved query performance. Parallelized query implementation was also used to improve performance.

## Supplementary Information

**Table S1.** Phenotype definitions and representative flow cytometry plots.

**Figure S1.** Histograms of 79 H2P2 traits. For flow cytometric data, raw phenotype values are used. For cytokine data, concentrations are log2 transformed.

**Figure S2.** Repeatability of 79 H2P2 traits. Repeatability was calculated as the inter-individual component of variance from ANOVA of 3 measurements of LCLs taken on sequential cell passages. Phenotypes are color-coded by stimuli. All traits have significant repeatability (p<0.05 marked by asterisk), except for three of the toxoplasma traits.

**Figure S3.** Minimal correlation between EBV copy number and H2P2 traits. The regression slopes of EBV copy number (from (*46*)) vs. the phenotypic values of each H2P2 trait (transformed into a Z-score and with population as a covariate) were determined. For all H2P2 traits grouped by biological category, slopes were rank ordered and plotted with the 95% confidence interval. Only two traits (IL12P70_Chlamydia and S_typhi_3_5hr_median_GFP) had slopes that were significantly non-zero at a nominal p<0.05 threshold, and even for these traits, EBV copy number accounted for only a small proportion of total variance (r^2^=0.028 and 0.025).

**Table S2.** Linear regression of EBV copy number and H2P2 traits.

**Figure S4.** Heritability of 79 H2P2 traits by parent-offspring regression. Individual plots for all H2P2 traits are shown with parent-offspring regression lines with (blue) and without (red) batch covariate. Estimated h^2^ and p-value for parent-offspring regression with batch covariate are listed on each plot.

**Table S3.** Heritability of 79 H2P2 traits by parent-offspring regression and SNP-based heritability.

**Figure S5.** QQ plots of 79 H2P2 traits. All SNPs with MAF > 5% are shown.

**Figure S6.** Association of rs2869462 with *CXCL10* mRNA. rs2869462 is associated with expression of the *CXCL10* mRNA (p=7×10^−7^) from the RNAseq data in (*57*)) in the same direction as the H2P2 *CXCL10* protein data.

**Table S4.** COLOC analysis of the CXCL10 region.

**Table S5.** eMERGE PheWAS results for 16 H2P2 hits. All eMERGE phenotypes with Bonferroni corrected p < 0.05 are shown.

